# Intranasal Administration of ACIS KEPTIDE™ Prevents SARS-CoV2-Induced Acute Toxicity in K18-hACE2 Humanized Mouse Model of COVID-19: A Mechanistic Insight for the Prophylactic Role of KEPTIDE™ in COVID-19

**DOI:** 10.1101/2020.11.13.378257

**Authors:** Gunnar Gottschalk, James F Keating, Kris Kesler, Konstance Knox, Avik Roy

## Abstract

Previously, we have demonstrated that ACIS KEPTIDE™, a chemically modified peptide, selectively binds to ACE-2 receptor and prevents the entry of SARS-CoV2 virions *in vitro* in primate kidney Cells. However, it is not known if ACIS KEPTIDE™ attenuates the entry of SARS-CoV2 virus *in vivo* in lung and kidney tissues, protects health, and prevent death once applied through intranasal route. In our current manuscript, we demonstrated that the intranasal administration of SARS-CoV2 (1*10^6^) strongly induced the expression of ACE-2, promoted the entry of virions into the lung and kidney cells, caused acute histopathological toxicities, and mortality (28%). Interestingly, thirty-minutes of pre-treatment with 50 μg/Kg Body weight ACIS normalized the expression of ACE-2 via receptor internalization, strongly mitigated that viral entry, and prevented mortality suggesting its prospect as a prophylactic therapy in the treatment of COVID-19. On the contrary, the peptide backbone of ACIS was unable to normalize the expression of ACE-2, failed to improve the health vital signs and histopathological abnormalities. In summary, our results suggest that ACIS is a potential vaccine-alternative, prophylactic agent that prevents entry of SARS-CoV2 *in vivo*, significantly improves respiratory health and also dramatically prevents acute mortality in K18-hACE2 humanized mice.

**Highlights:**

- ACIS KEPTIDE stimulates the internalization of ACE-2 receptor (Fig. 2) and buffers the membrane localization of ACE-2 receptors (Fig. 2, 6 & 8). Intranasal inoculation of SARS-CoV2 upregulates the expression of ACE-2 in lung epithelium (Fig.6) and kidney tubular cells (Fig.8). ACIS KEPTIDE normalizes the expression of ACE-2 in the kidney tubular cells of virus-treated K18-hACE2mice (Fig. 8).
- ACIS KEPTIDE™ <u>completely prevents</u> the entry of SARS-CoV2 in Bronchiolar epithelium (Fig.6), alveolar parenchyma (Fig. 6), and kidney tubular cells (Fig.8).
- ACIS KEPTIDE™ improves the pulmonary (Fig. 5) and renal pathological changes (Fig. 7) caused by the SARS-CoV2 virus insult.
- Intranasal administration of 0.05% Beta-propiolactone (βPL)-inactivated SARS-CoV2 (1 *10^6^) causes significant death (28%) in K18-hACE2 humanized mice after 24 hrs of intranasal inoculation (**<u>Supplemental videos</u>**) suggesting that SARS-CoV2 does **<u>not</u>** require its infective properties and genetic mechanism to be functional to cause mortality.
- The peptide backbone of ACIS KEPTIDE™ provides much less and insignificant protection in the prevention of pathological changes in Lungs (Fig.5 & 6) and Kidney (Fig.7 & 8). Peptide failed to normalize the upscaled expression of ACE-2 in kidney tubular cells (Fig.8) of SARS-CoV2-treated K18-hACE2 mice.

## Introduction

COVID-19, a severe acute respiratory disease [1, 2], is primarily caused by a viral strain named as Coronavirus-2 (SARS-CoV2) [3]. As an entry mechanism of the virus into host cell, A recent study [4] has demonstrated that SARS-CoV2 employs its membrane-bound “S-glycoprotein” to attach with the host receptor ACE-2. Although, other mechanisms such as attachment of virus to the polysaccharide molecule heparan sulphate [5] in the cell membrane also facilitates the infectivity of the virus, until now, the interaction of SARS-CoV2 with ACE-2 protein has been considered as a primary mechanism of viral entry [6] [7].

In our previous study [8], we reported that ACIS KEPTIDE™, a chemically modified peptide with a strong affinity towards ACE-2 receptor, efficiently nullified the interaction between ACE-2 and S-glycoprotein of SARS-CoV2. As a result, the entry of the SARS-CoV2 virions into host cells had been severely impaired. Based on the results derived from different experiments such as cytopathic effect assay, plaque assay and dual immunofluorescence analyses, we concluded that ACIS KEPTIDE™ indeed inhibited the attachment and entry of SARS-CoV2 virions in VEROE6 primate kidney cells. Moreover, our results also demonstrated that the chronic administration of KEPTIDE through intranasal pathway did not cause the loss of body weight, changes in vitals including body temperature, oxygen saturation, and heart rate nullifying any health-related toxicity due to KEPTIDE-treatment. Although that experiment was performed in aged BALB/C mice, understanding the toxicity of KEPTIDE is warranted in better animal model that express human ACE-2 receptor.

Therefore, in our current work, we included 6-8 weeks old K18-hACE2 mice. This strain is a humanized strain that express human ACE-2 receptor under the guidance of K18 promoter. K18-hACE2 mice display an acute lung and kidney injuries resulting significant mortality upon inoculation with SARS-CoV2 [9-11]. Therefore, to avoid significant mortality in these mice and for the safety of users, we inactivated virus with 0.05% beta-propiolactone (βPL). Although the treatment with 0.05% β-propiolactone has been reported to substantially reduce the infectivity of the virus, the ultrastructure of the external capsid remains unaltered and its antigenicity is also preserved [12, 13] that allows us to perform the entry experiment of SARS-CoV2 in lung tissue. To explore the effect of ACIS on preventing the entry of virions, we performed an *in vivo* experiment with the intranasal treatment of ACIS for 30 minutes followed by the intranasal inoculation of SARS-CoV2 for another 24 hrs. Next, we performed an array of experiments in lungs and kidney of these humanized animals to confirm the role of ACIS KEPTIDE in the entry of virions through ACE-2 receptor, the regulation of ACE-2 receptor, amelioration of acute histopathological changes, and mortality.

## Results

### Exploring toxicity in K18-hACE2 animals after chronic intranasal administration of ACIS KEPTIDE™

Eight to 10 weeks old K18-hACE2 mice (*n=8*; 4 males and 4 females) were intranasally administered with 50 μg/Kg Bwt KEPTIDE every day for 10 days as described under method section. Average baseline body weights were 18.25±1.4 gms and 15.65± 2.3 gms for males and females, respectively. Each day animals were recorded for body weight (Fig. 1A), body temperature (Fig. 1B), heart rate (Fig. 1C), and oxygen saturation (Fig. 1D) as described previously [8]. As evident from figure 1A, both male and female mice maintained their body weights throughout the study. On average, there was a decrease in body weight (14.59 ± 2.32) recorded in female mice on day 9 that happened most likely due to the loss of water-weight. However, that drop was not statistically significant while comparing with the baseline value (p>0.05; =0.2015). On contrary, male mice did not display any loss in body weights. Instead, we observed consistent increase in body weights from day 6. Again that increase was not statistically significant once compared with the baseline value (p>0.05; =0.0762). Daily administration of KEPTIDE did not alter body temperature (Fig. 1B), heart rate (Fig. 1C), and oxygen saturation (Fig. 1D). There was only partial and non-significant elevation of blood pressure on day 9 (baseline value of 448± 12.5 to 489± 10.7) in all females, otherwise all vital parameters were found to be stable. No mortality was observed across the study in any animal.

**Fig. 1.**
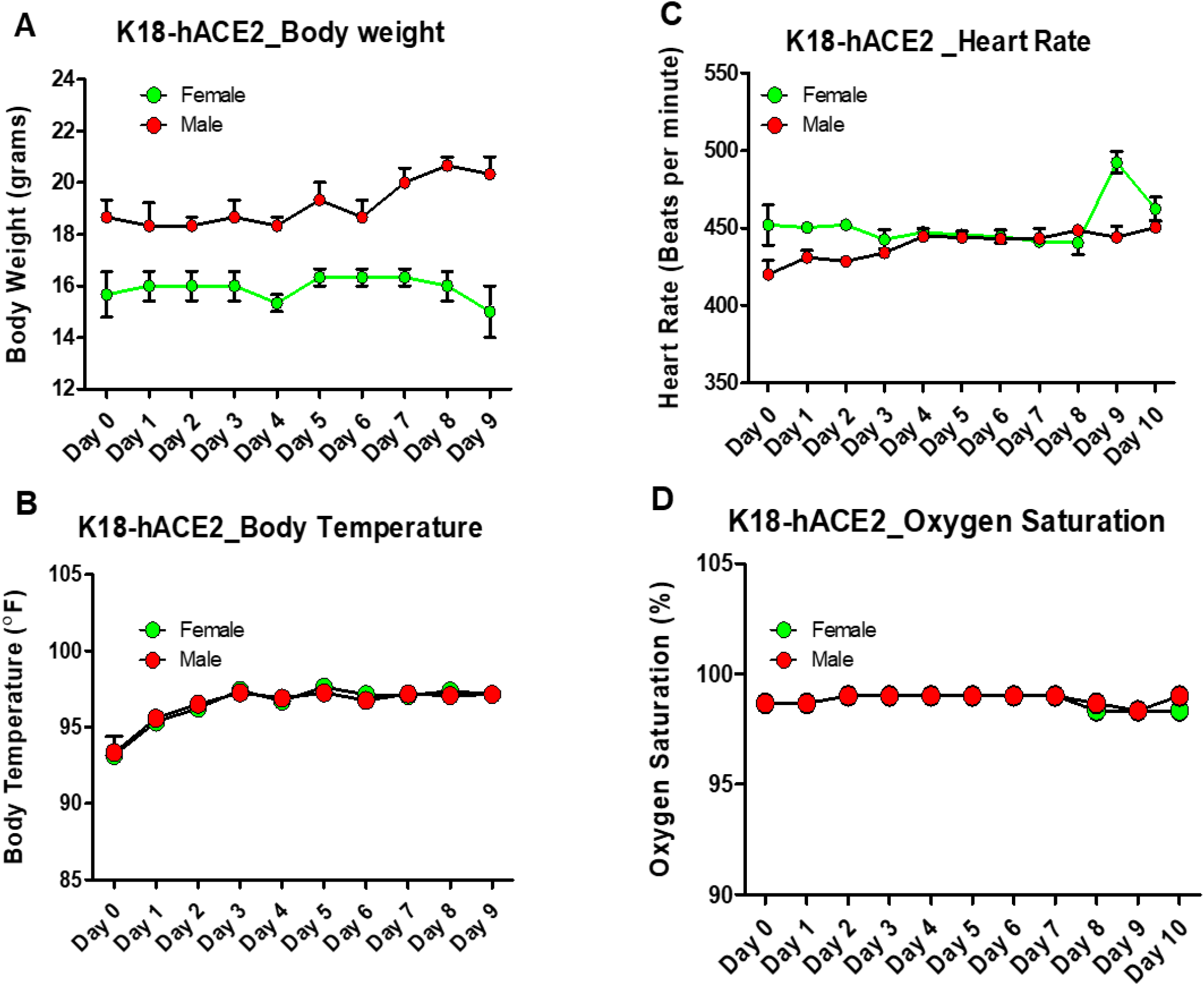
The effect of intranasally administered of ACIS KEPTIDE on the health parameters of K18-hACE2 mice. (A) Eight-to-ten weeks old K18-hACE2 mice were administered intranasally with ACIS KEPTIDE at a dose of 50 μg/Kg bodyweight for 10 days (n=8 per group; 4 males and 4 females). Different health parameters sch as (A) body temperature, (B) body weight, (C) heart rate, and (D) oxygen saturation were monitored starting from day 0 to day 10. Results are mean ± SEM of three different experiments. Significance of mean between two groups were tested with paired t-test (p = 0.234 for body temperature; p = 0.00109841 for body weight; p = 0.0981 for heart rate and p = 0.7121 for oxygen saturation). Results are mean ± SEM of three independent experiments.

### ACIS KEPTIDE™ prevented the mortality and protected health of SARS-CoV2-infected K18-hACE2 mice

Since ACIS did not display any mortality and health-related issue in K18-hACE2 mice, next we wanted to study if ACIS could also prevent the death and improve health-associated toxicities in these mice upon exposure to SARS-CoV2 virions. Wuhan-standard SARS-CoV2 is highly infectious and therefore complete inactivation of virus is required in order to minimize the health risk associated with the administration procedure, perfusion, surgery, post-surgical processing of tissue, and histological analyses. Accordingly, the inactivation was performed with incubating viral stock with 0.05% βPL as described elsewhere [13] and also mentioned under method section. Based on our plaque assay (Supplementary Fig. 1), we determined that 0.05% βPL indeed significantly inactivated the SARS-CoV2 virus.

In order to assess the protective role of ACIS KEPTIDE in the entry of SARS-CoV2 in lung epithelium, 8-10 weeks old K18-hACE2 mice (*n=8* per group; 4 males and 4 females) was inoculated with 50 μg/Kg bwt ACIS intranasally for 30 minutes followed by the administration inactivated 1×10^6^ SARS-CoV2 virus particles. Thirty minutes of pre-incubation with ACIS was justified based on our result of immunofluorescence analyses performed in vitro in human lung epithelial cells Calu-3. We observed that thirty minutes of exposure to 25 μM of ACIS strongly induced the internalization of ACE-2 receptor in human Calu-3 lung cells and continued to downregulate the *de novo* expression of ACE-2 receptor (Fig. 2) for 1, 2 and 6 hours of incubation. Based on that result, we decided to pre-treat the mice (n=8 per group) with 50 μg/Kg Bwt KEPTIDE for 30 mins followed by the treatment with 1 *10^6^ virus for 24 hrs. After another 24 hrs, mice were analyzed for the vital signs of health and mortality. <u>After 24 hrs</u>, we observed following.

**Fig. 2.**
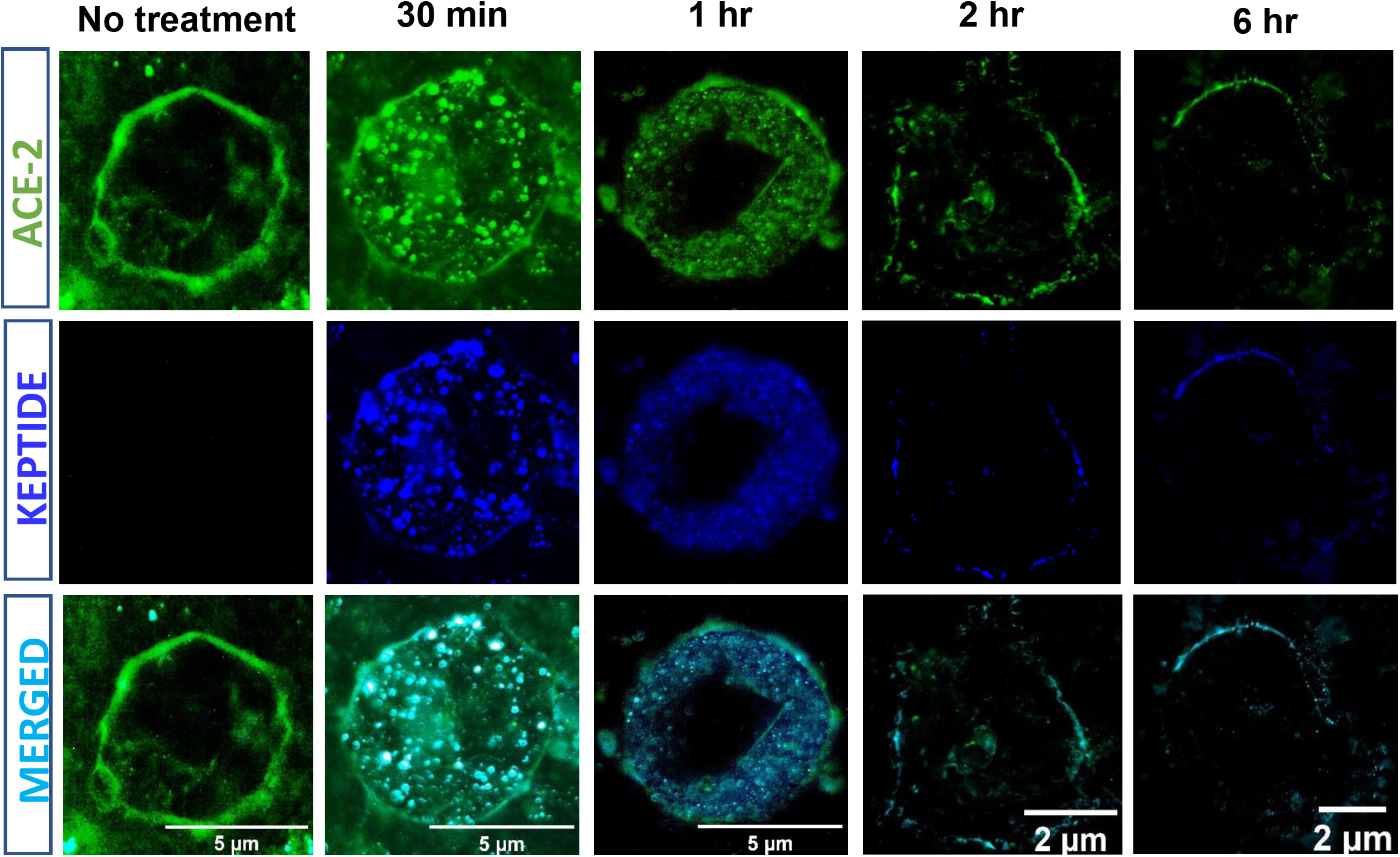
Effect of KEPTIDE on the Expression of ACE-2 Receptor on the Membrane of CALU-3 Human Lung Cells. Human lung epithelial cells. CALU-3 cells were grown in complete DMEM cells for 2 days until it reached 70% confluency followed by starving with serum for 2 hrs. After that, 25 μM of ACIS KEPTIDE were treated for 30 mins, 1 hr, 2 hrs and 6 hrs. After each time point cells were fixed and stained for ACE-2 (Green; Rabbit anti-ACE-2 antibody; Abcam; 1:250 dilution) and KEPTIDE (blue). Thirty minutes of KEPTIDE treatment significantly stimulated the internalization of ACE-2 along with KEPTIDE. Subsequent incubation periods displayed significant down-regulation of ACE-2 receptors. Experiments were confirmed after three different experiments.

*First*, there was significant mortality (28%; 2 out of 7 animals) in virus-treated animals. Two females (66%; 2 out of 3 females) died (Supplementary video 1). *Second*, virus-treatment caused significant loss of body temperature (Fig. 3A). *Third*, another striking feature is abrupt loss of body weight (Fig.3D). One male showed 8 gm loss of body weight (26 gm day 0 to 18 gm day post inoculation; tail mark1 male). Otherwise, across the virus-treatment group there was average 3.04 ± 2.54 gms of body weight loss (Fig. 3D). However, there was no statistical significance (p>0.05) while measuring significance of mean between groups with paired t test as because the sample size changed due to mortality. All virus-treated mice are moribund, with low body temperature (Fig. 3A), abnormally low heart rate (Fig. 4A; ^*\*\*\**^*p*<0.0001), severe respiratory stress (Fig. 4D; O_2_ saturation less than 95%; ^*\*\*\**^*p*<0.005), and motor impairment. All mice displayed severe physical discomfort as evident with hunched posture, scruffy skin, frequent shivering, and hindlimb dragging (Supplemental video 1).

**Fig. 3.**
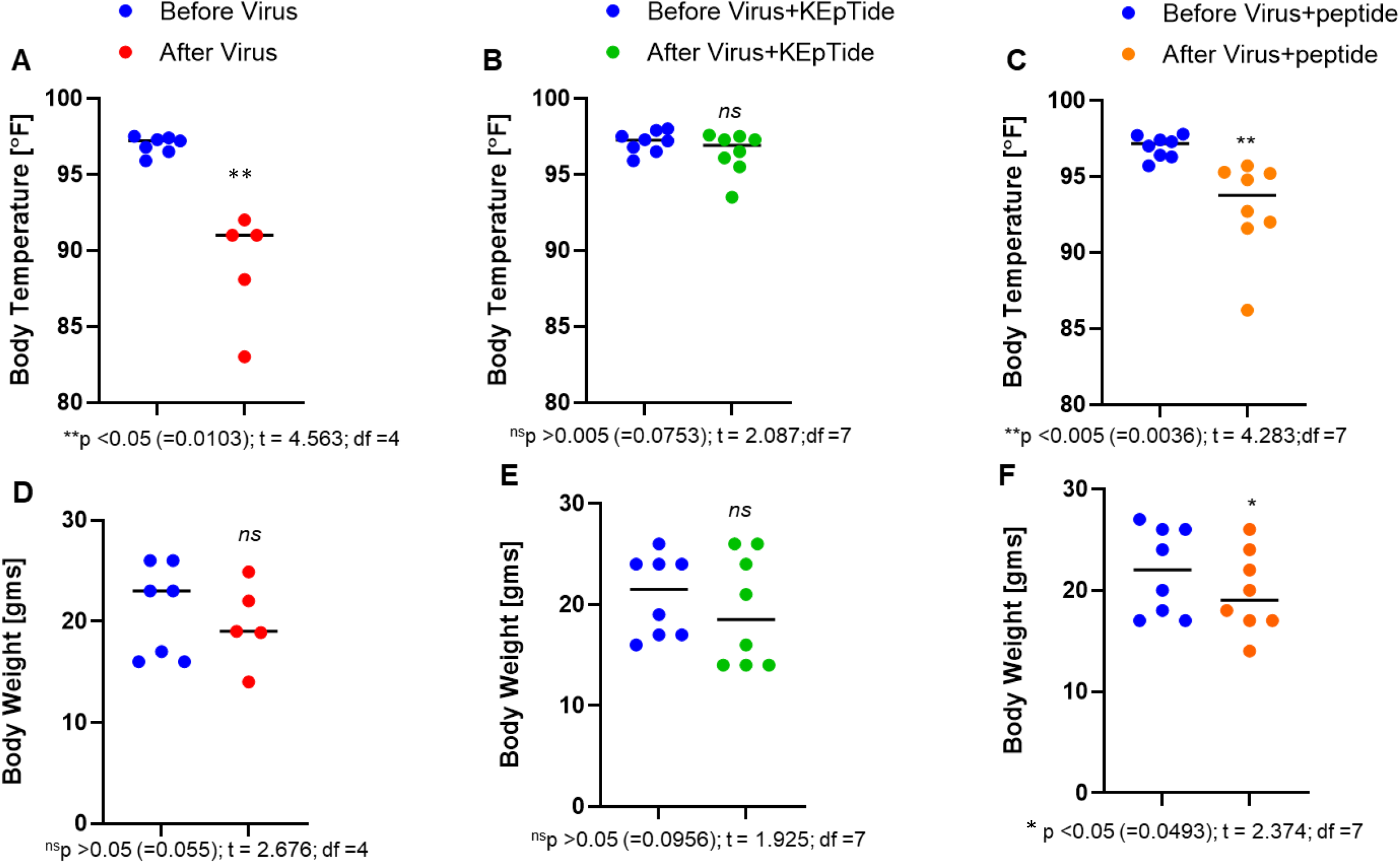
Effect of KEPTIDE on the Regulation of Body Temperature and Body Weight in K18-hACE-2 Humanized Mice Infected with SARS-CoV2 Virus for 24 hrs. Eight to ten weeks old K18-hACE2 mice were intranasally administered with 50 μg/Kg Bwt KEPTIDE or 50 μg/Kg Bwt peptide for 30 mins followed by inoculation with 1*10^6^ inactivated virus. Virus was inactivated with 0.05% β-propiolactone as mentioned under method section. In group1 mice (n=7) were inoculated with virus only; in group 2, mice (n=8) were treated with virus +KEPTIDE; and, in group 3, mice (n=8) were treated with virus +peptide. After 24 hrs, significant mortality (2 females of 7 mice) was observed in group 1, but not other groups. (A-C) Body temperature was monitored. Both virus only- and virus+peptide-treated animals have significantly low body temperature once compared with the baseline body temperature recorded one day before the procedure. (D-F) Body weight was monitored in three groups one day before and after of procedure. No significant change in body weight observed in group1 and 2, however peptide treatment could not protect the loss of body weight due to virus inoculation. Mortality is the possible confounder in group1 while measuring body weight. Results are mean ± SEM of 7-8 animals. Related p value with descriptive t statistics were mentioned below each histogram.

**Fig. 4.**
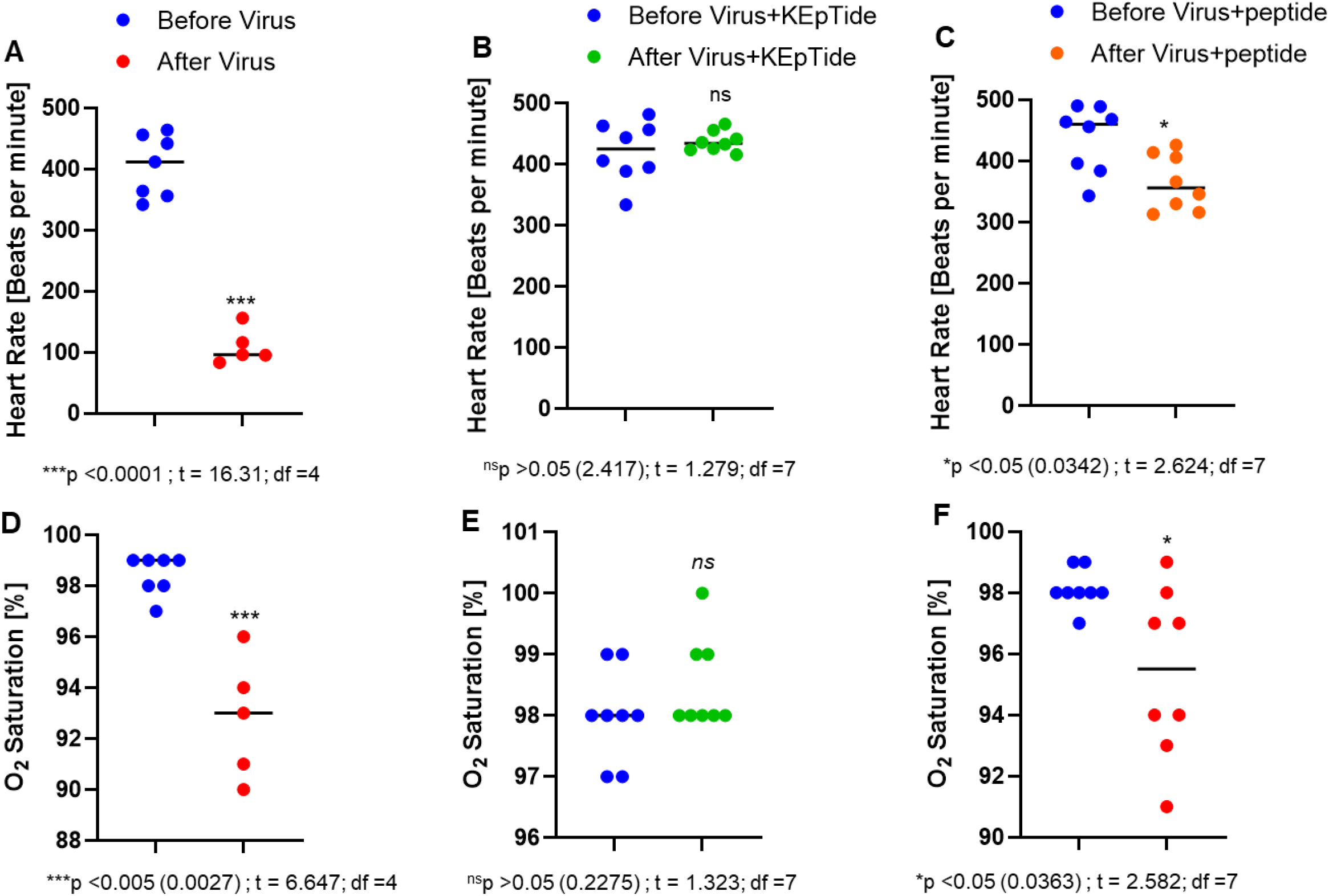
Effect of KEPTIDE on the Regulation of Heart Rate and O2 saturation in K18-hACE-2 Humanized Mice Infected with SARS-CoV2 Virus for 24 hrs. Eight to ten weeks old K18-hACE2 mice were intranasally administered with 50 μg/Kg Bwt KEPTIDE or 50 μg/Kg Bwt peptide for 30 mins followed by inoculation with βPL-inactivated 1*10^6^ virus. In group1 mice (n=7) were inoculated with virus only; in group 2, mice (n=8) were treated with virus + KEPTIDE; and, in group 3, mice (n=8) were treated with virus + peptide. (**A-C**) Body temperature was monitored after 24 hrs. Both virus only- and virus+peptide-treated animals have significantly low heart rate once compared with the baseline heart rate recorded one day before the procedure. (**D-F**) Oxygen (O_2_) saturation was monitored in three groups one day before and after of procedure. Significant change in O_2_ saturation observed in group1 and 3. Interestingly, KEPTIDE treatment protected the respiratory health even after virus inoculation. Results are mean ± SEM of 7-8 animals. Related p value with descriptive t statistics were mentioned below each histogram.

Interestingly, KEPTIDE-treated and virus-infected mice displayed no mortality (Supplementary video 2), normal body temperature (Fig. 3B), body weight (Fig. 3E), heart rate (Fig.4B), and Oxygen saturation (Fig. 4E). Skin tone is normal with shiny fur. Mice did not show any motor impairment and no physical discomfort such as frequent shivering, curved posture and hind limb dragging. Male and female mice dis not display any gender biasness in terms of health and mortality. There was no change in body weight.

In contrast, peptide-(same amino acid sequence backbone, but no Knock-End tagging and neutralization) treated & virus-treated mice (n=8) did not display any mortality (Supplementary video 3) but significant impairment in movement (2 out of 4 male and 2 out of 4 female) as indicated with hunchback posture, frequent shivering, scruffy skin and dizziness. These mice display decreased body temperature (Fig. 3C; ^\**\**^*p<0*.*005*), body weight (Fig. 3F; ^*\**^*p<0*.*05*) low heart rate (Fig. 4C; ^*\**^*p<0*.*05*), and severe respiratory stress (Fig. 4F; ^*\**^*p<0*.*05*). Taken together our results suggest that even after significant attenuation of infectivity, βPL-treated SARS-CoV2 virus is still able to cause acute health issues and mortality.

### ACIS KEPTIDE prevented SARS-CoV2-induced acute histopathological changes in lungs and kidneys of K18-hACE2 mice

Since intranasal inoculation of βPL-inactivated virus caused significant health toxicity and pre-treatment of ACIS KEPTIDE, but not peptide, dramatically prevented the health issues, next we wanted to analyze the histopathological abnormalities in lungs and kidneys in virus-treated animals. We were also interested to analyze if ACIS KEPTIDE prevented these abnormalities. Therefore, we first performed hematoxylin & eosin (H&E) staining in paraffin-embedded lungs and kidney sections of vehicle-, virus-, virus +KEPTIDE, and virus+peptide-treated groups. A detailed analysis in H&E-stained slides demonstrated that intranasal inoculation of βPL-treated virus, but not βPL-treated uninfected VERO sup (vehicle; Fig. 5A), caused the swelling of alveolar parenchyma with increased number of cells (Fig. 5B). Interestingly, thirty-minutes pre-treatment of 50 μg/Kg Bwt KEPTIDE (Fig. 5C), but not its peptide backbone (Fig. 5D), significantly protected the alveolar space, thickness of the alveolar wall and cellularity of alveolar parenchyma. Further analyses of bronchiolar epithelium revealed that virus treatment (Fig. 5F & Fii), but not vehicle (Fig. 5E & Ei), significantly damaged the integrity of the epithelial membrane with hemorrhagic response in the surrounding pulmonary blood vessels. On the other hand KEPTIDE pre-treatment not only protected the integrity of the bronchiolar epithelium (Fig. 5G & Giii), but also prevented hemorrhagic response. Interestingly, peptide treatment could not block the virus-mediated damage of bronchiolar epithelium (Fig. 5H & Hiv) and pulmonary hemorrhagia.

**Fig. 5.**
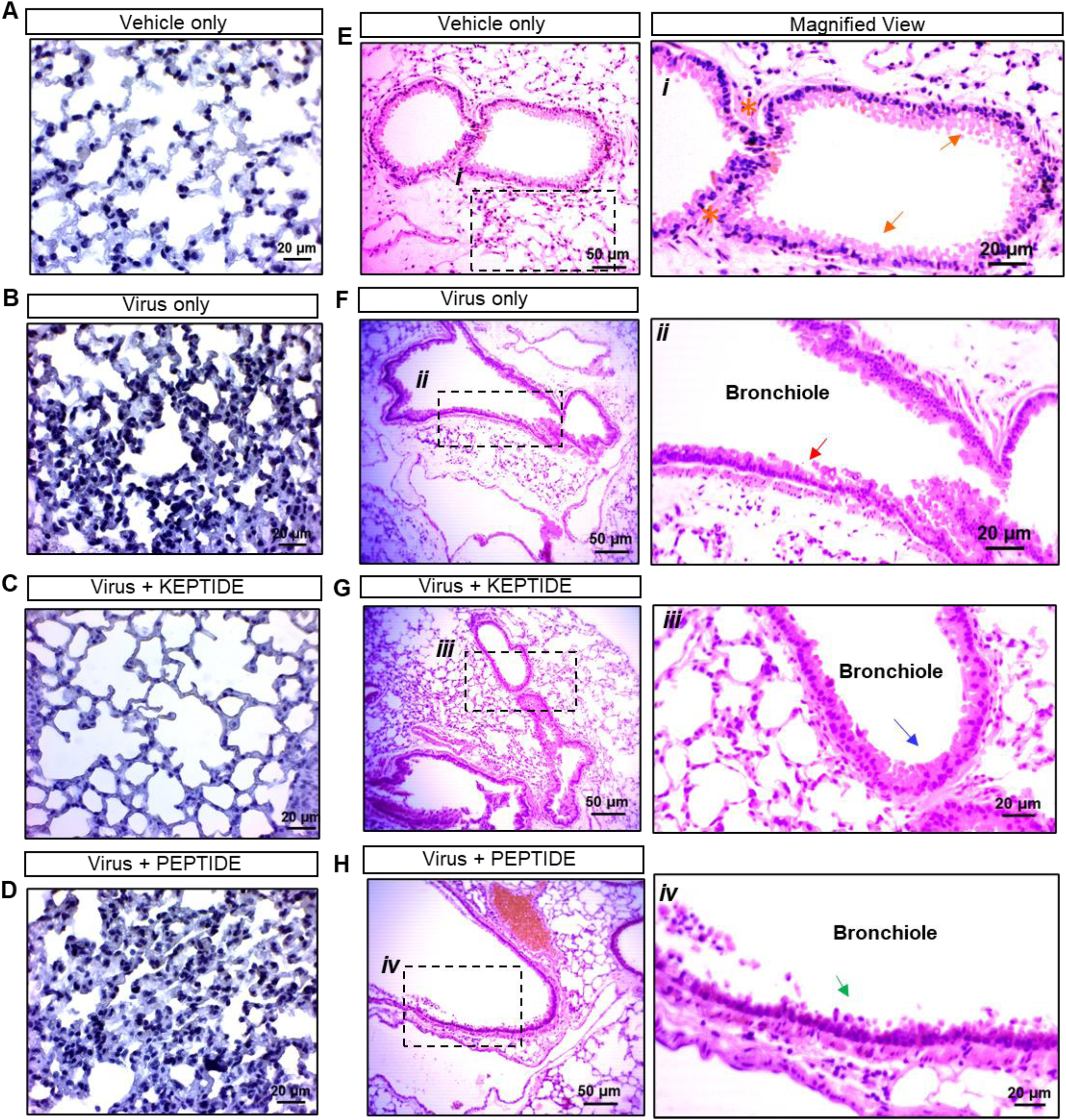
ACIS KEPTIDE protects acute histopathological changes in Lungs of K18-hACE2 mice twenty-four hours post SARS-CoV2 inoculation. Eight to ten weeks old K18-hACE2 mice (n=7-8 per group) were intranasally administered with 50 μg/kg Bwt KEPTIDE or 50 μg /kg Bwt peptide for 30 mins followed by inoculation with βPL-inactivated 1*10^6^ virus. In group1 mice (n=8) were inoculated with vehicle only (0.05% βPL-treated media); in group 2, mice (n=7 were treated with virus only (0.05% βPL-inactivated; in group 3, mice (n=8) were treated with virus +KEPTIDE; and, in group 4 mice (n=8) were treated with virus + PEPTIDE. (**A-D**) Hematoxylin Background staining of small airway alveolar parenchyma. (**E-H**) H & E staining of Bronchiolar epithelium, surrounding cartilaginous and alveolar parenchyma. Magnified views of bronchiolar epithelia of (**Ei**) Control (orange arrow indicates intact epithelial lining: orange star demonstrates preserved connective tissue), (**Fii**) Virus only (thin red arrow indicates the degenerated bronchiolar epithelium), (**Giii**) Virus + KEPTIDE (thin blue arrow indicates protected epithelium), and (**Hiv**) Virus + Peptide-treated (thin green arrow indicates degenerated epithelium). Results are confirmed after three different experiments in 7-8 animals.

Is this KEPTIDE-mediated protection of acute toxicity due to the prevention of viral entry in lung epithelium? To address this concern, we performed a dual immunohistochemical analyses of ACE2 and Spike glycoprotein (Fig. 6). Surprisingly, twenty-four hour treatment with SARS-CoV2 significantly stimulated the expression of ACE-2 in the outer (Fig. 6 B*c*) and inner layers (Fig.6 B*d*) of bronchiolar epithelium compared to vehicle treatment (Fig, 6A*a* & 6 A*b*).This upregulation of ACE-2 expression by SARS-CoV2 was unexpected and might facilitate the entry of virions through lung epithelium [14]. Thirty-minutes pretreatment with ACIS KEPTIDE (Fig. 6C*e* & 6C*f*), but not peptide (Fig. 6D*g* & 6D*h*), strongly downregulated the expression of ACE-2 and normalized its expression to the basal level. Accordingly, we observed the infiltration of SARS-CoV2 in the bronchiolar parenchyma (brown arrow) in virus- and virus + peptide-treated groups, whereas KETIDE treatment significantly protected the entry of virions (indicated with brown arrow). The result was corroborated with the quantification of numbers of ACE-2-immunoreactive (*ir*) cells in bronchiolar epithelium in all four groups (Supplementary Fig. 2A). To further evaluate if these brown signals were not artifacts, we confirmed the size of these particles in reference to eosin-stained nuclei. However, these viral particles were inactivated and therefore lost the proliferative properties. As a result, the load of virions was not overwhelming in lung parenchyma.

**Fig. 6.**
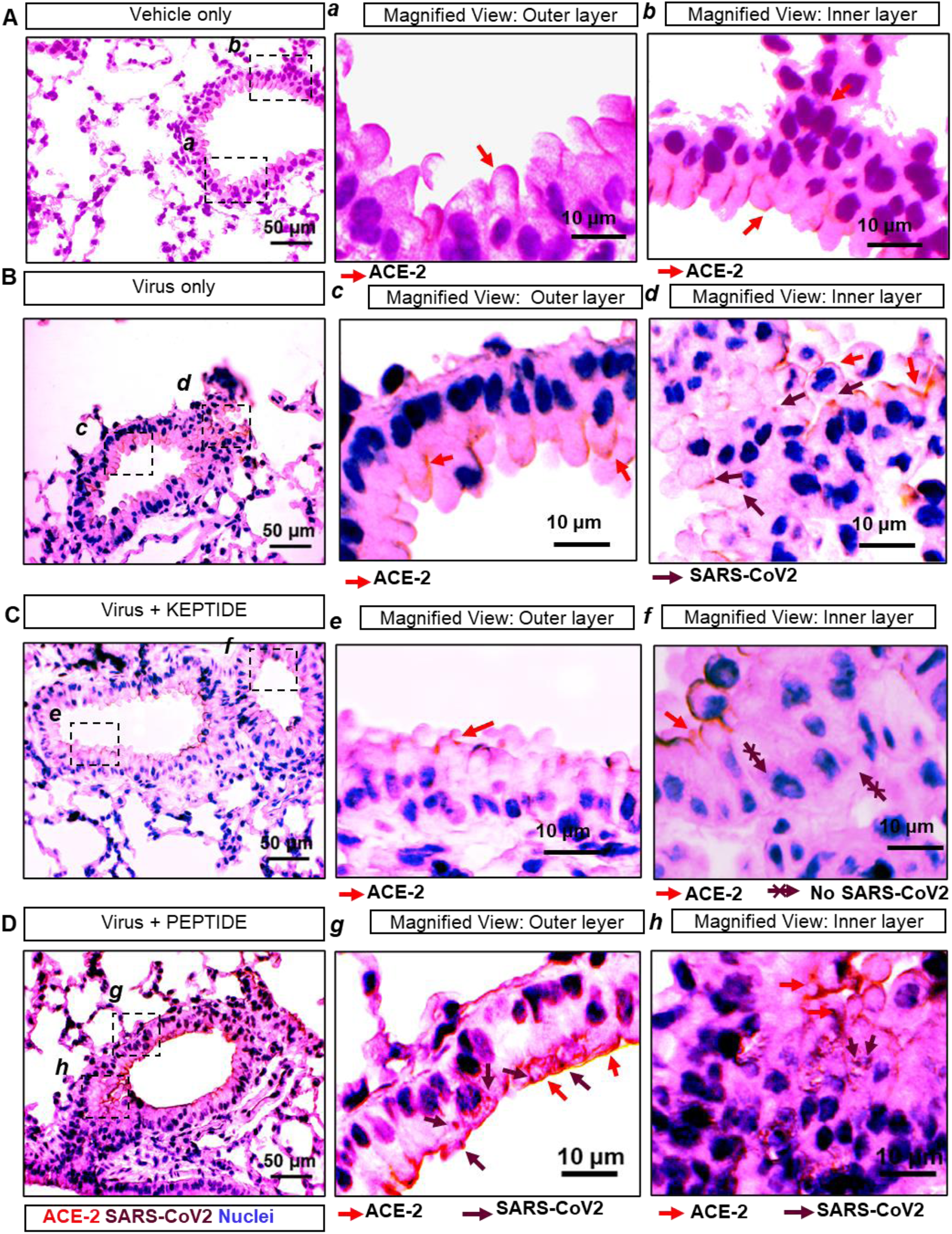
Protective effect of ACIS KEPTIDE on the expression of ACE-2 and the entry of SARS-CoV2 virions in lung of SARS-CoV2-insulted K18-hACE2 mice. (**A-D**) Dual IHC staining of ACE-2 (red) and SARS-CoV2 (brown) in bronchiolar epithelium in (**A**) vehicle-treated (0.05% βPL-inactivated VEROE6 sup, (**B**)virus, (**C**) virus + KEPTIDE, and (**D**) virus + peptide-treated K18-hACE2 mice (n= 7-8). (***a-h***) Magnified views of outer layers and inner layers of bronchiolar epithelia of respective images enclosed in a dotted squares. Arrows were justified in the bottom of each image. Results are confirmed after three independent experiments.

### ACIS KEPTIDE™ prevents the entry of SARS-CoV2 in the Kidney Tubular Cells

According to a recent report [15], tubular cells of Kidney strongly express ACE-2 receptors. Next, we explored pre-treatment of ACIS was able to nullify the entry of SARS-CoV2 virions in renal tubular cells. We observed that SARS-CoV2 ACIS (Fig. 7A), but not vehicle (Fig. 7B), significantly altered the integrity of kidney cortex with the formation of vacuoles around glomeruli, whereas ACIS KEPTIDE protected the integrity of kidney cortex (Fig. 7C) with unaltered morphology of glomeruli, proximal and distal convoluted tubular cells. Surprisingly, the kidney cortices of the peptide backbone-treated group displayed a hemorrhagic response (Fig. 7D). However, we did not observe any vacuolization in the kidney cortex of virus +peptide-treated group. The result was further corroborated with the quantification data (Supplemental figure 2B). Next, we monitored the regulation of ACE-2 receptor in kidney tubular cells and also evaluated how that expression could alter the entry of virions in these cells. Interestingly, our dual IHC analyses revealed that overnight inoculation of SARS-CoV2 virus indeed upregulated the expression of ACE-2 in tubular epithelia (Fig. 8A & 8B; Supplementary Fig. 2B), whereas pre-treatment of KEPTIDE (Fig. 8C), but not peptide (Fig. 8D), normalized the expression of ACE-2 (Supplementary Fig. 2B). In addition to that, we also observed that pre-treatment with KEPTIDE significantly inhibited the entry of SARS-CoV2-promoted entry of virions in kidney tubular cells and also through Bowman’s capsule of glomerulus.

**Fig. 7.**
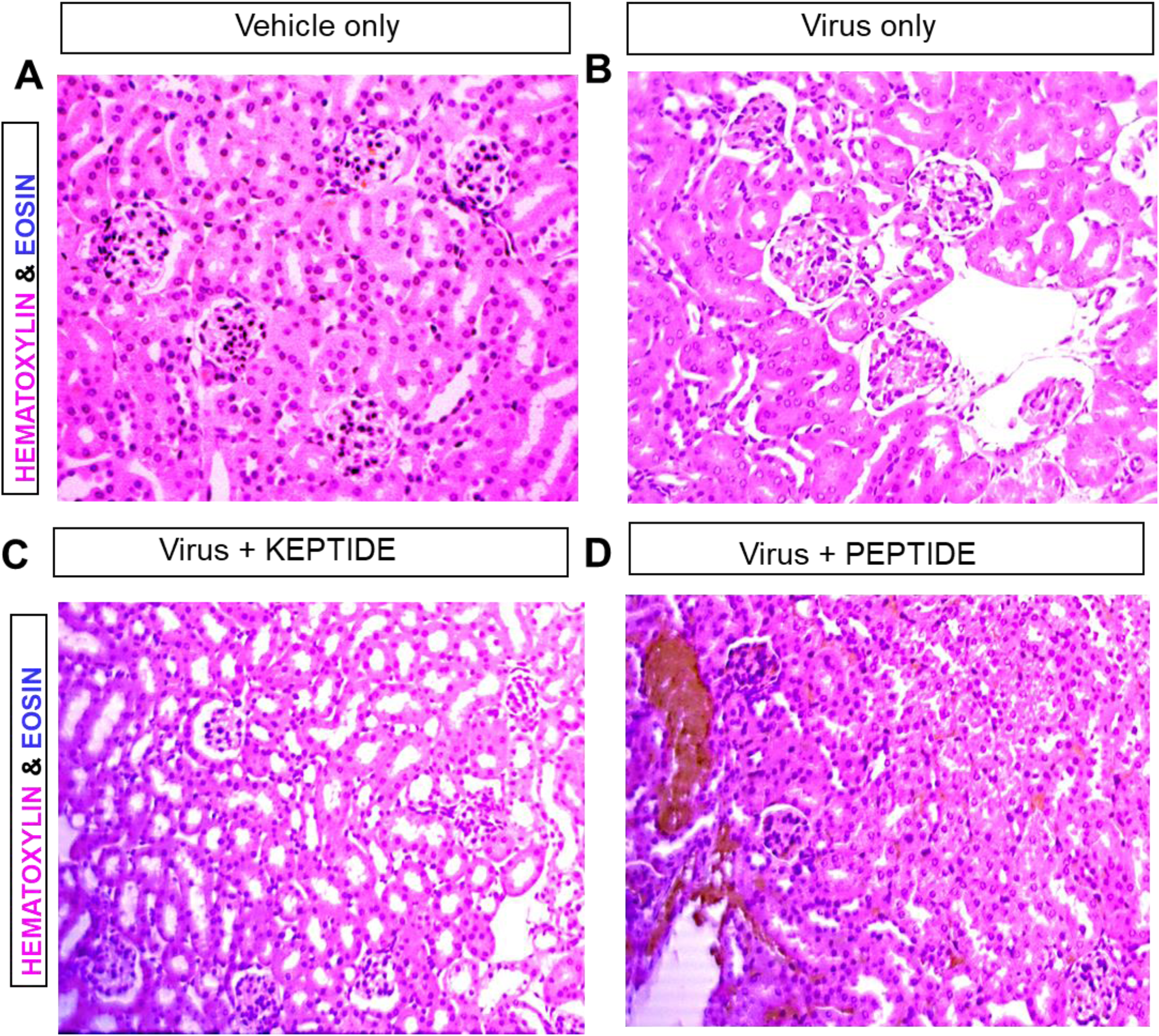
Effect of ACIS KEPTIDE on the protection of acute histopathological changes in Kidney of K18-hACE2 mice twenty-four hours post SARS-CoV2 inoculation. Eight to ten weeks old K18-hACE2 mice (n=7-8 per group) were intranasally administered with 50 μg/kg Bwt KEPTIDE or 50 μg/kg Bwt peptide for 30 mins followed by inoculation with βPL-inactivated 1*10^6^ virus. In group1 mice (n=8) were inoculated with vehicle only (0.05% βPL-treated VEROE6 sup); in group 2, mice (n=7) were treated with virus only (0.05% βPL-inactivated; in group 3, mice (n=8) were treated with virus +KEPTIDE; and, in group 4 mice (n=8) were treated with virus + PEPTIDE. (**A-D**) H & E staining of kidney cortex with detailed structures of glomeruli, tubular epithelium, and Bowman’s capsule in all 4 groups. Results are confirmed after three independent experiments.

**Fig. 8.**
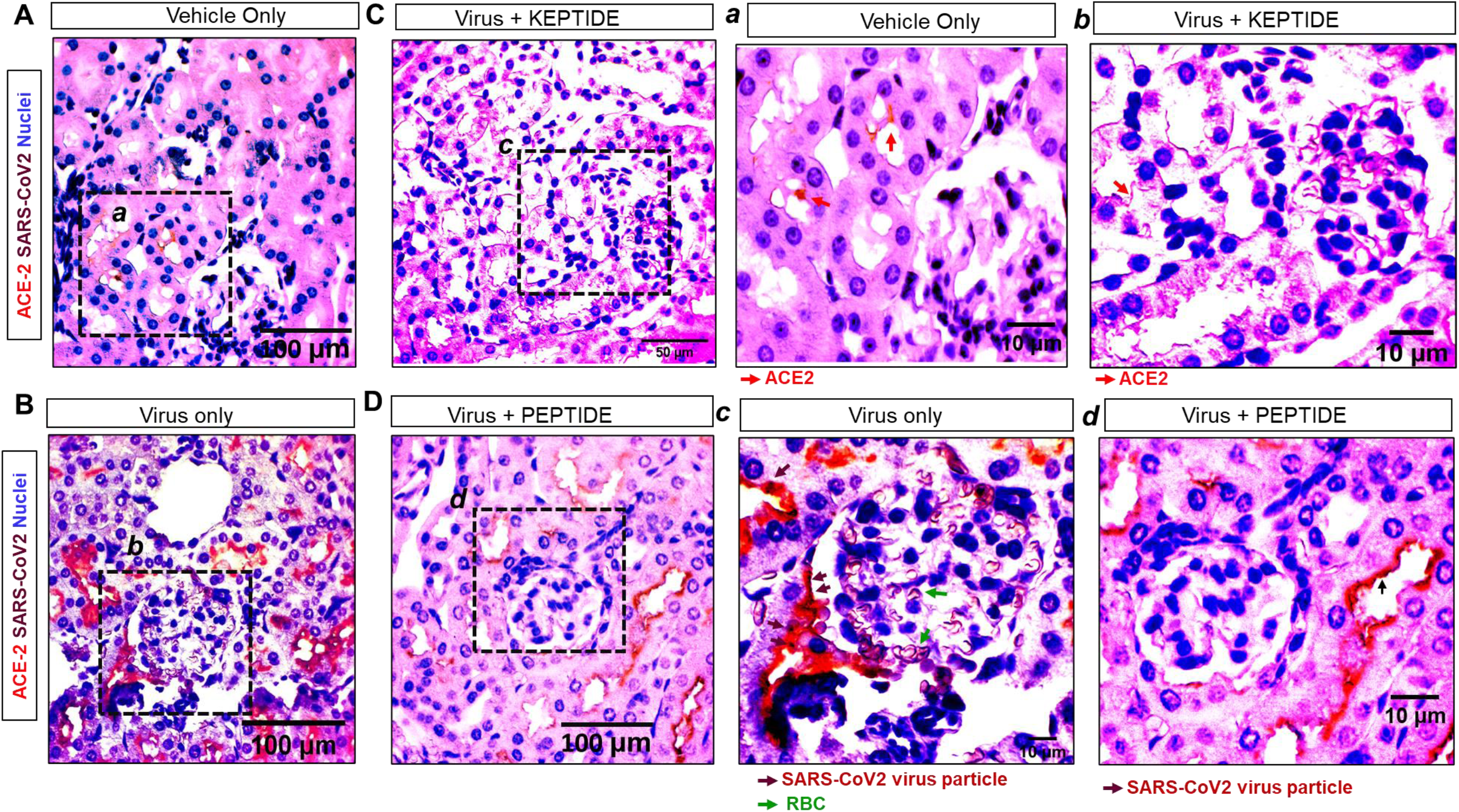
Protective effect of ACIS KEPTIDE on the expression of ACE-2 and prevention of SARS-CoV2 entry in kidney of K18-hACE2 mice twenty-four hours post SARS-CoV2 inoculation. Eight to ten weeks old K18-hACE2 mice (n=7-8 per group) were intranasally administered with 50 μg/kg Bwt KEPTIDE or 50 μg/kg Bwt peptide for 30 mins followed by inoculation with βPL-inactivated 1*106 virus. In group1 mice (n=8) were inoculated with vehicle only (0.05% βPL-treated VEROE6 sup); in group 2, mice (n=7) were treated with virus only (0.05% βPL-inactivated; in group 3, mice (n=8) were treated with virus +KEPTIDE; and, in group 4 mice (n=8) were treated with virus + PEPTIDE. (Results are confirmed after three independent experiments. (**A-D**) Dual IHC of ACE-2 (red) and SARS-CoV2 (brown) in tubular epithelium in (**A**) vehicle, (**B**) Virus, (**C**) Virus + KEPTIDE-, and (**D**) Virus + peptide-treated groups. (***a***) showed magnified view of kidney cortex vehicle-treated mouse with basal expression of ACE-2 (red), (***b***) magnified view of kidney of virus-tread animal. Upscaled expression of ACE-2(red) with degenerated Bowman’s capsule and invasion of SARS CoV2 (Brown arrow). (***c***) Magnified view of Glomerulus of Virus + KEPTIDE-treated group. Significantly less ACE-2 expression (red arrow) and no virus-infiltration were noted. (***d***) Elevated expression of ACE-2 in virus +Peptide-treated group. Results were confirmed after three different experiments in 7-8 animals.

## Discussion

### ACIS KEPTIDE stimulates the internalization and downregulation of ACE-2 receptor in Lung and Kidney epithelial cells

Bronchiolar epithelium is composed of tightly attached epithelial cells that are known to express wide-range of pattern recognition receptors dedicated to innate immune response for providing defense against invading microbes [16]. Apart from immune protection against microbes, these epithelial cells also strongly express ACE-2 [17], which is known for its vasodilative action. Recent studies indicate that ACE-2 is a selective receptor for the entry of SARS-CoV2 virions in lung epithelial cells. Upon binding to ACE-2, SARS-CoV2 virus gets internalized to the cytosol of the epithelial cell to initiate the early pathogenic events [17, 18]. ACIS KEPTIDE was designed to selectively inhibit the binding of S-glycoprotein of SARS-CoV2 with ACE2 and subsequently block the entry of SARS-CoV2 through ACE-2 receptor [8]. However, the effect of ACIS on the regulation of ACE-2 receptor is not known. In our present manuscript, we explored the effect of ACIS on the expression and turn-over of ACE-2 receptors in lung and kidney epithelial cells. *First*, our immunocytochemical analyses revealed that upon binding, ACIS stimulated the internalization of ACE-2 receptor *in vitro* in cultured human lung epithelial cells Calu-3. The effect was observed as early as 30 minutes treatment of 25 μM of KEPTIDE resulting the consistent downregulation of ACE-2 expression over longer period of incubation. *Second*, our dual IHC analyses indicated that SARS-CoV2 strongly induced the expression of ACE-2 in bronchiolar epithelial cells and the thirty minutes pre-treatment of ACIS efficiently normalized the virus-stimulated expression of ACE-2 *in vivo* in bronchiolar epithelial cells. *Third*, similar treatment of ACIS KEPTIDE also dramatically inhibited the expression of ACE 2 *in vivo* in renal tubular cells of K18-hACE2 mice. Fourth, pre-treatment with peptide backbone of ACIS was unable to restore the expression of ACE-2 in virus-inoculated mice. Combing all results, we confirmed that ACIS is a selective ACE-2 inhibitor that binds, internalizes, and balances the expression of ACE-2 preventing the entry of SAR-CoV2 in the host cell.

### ACIS KEPTIDE improves the pathological outcomes of SARS-CoV2-infected lung and kidney tissue

Intranasal inoculation of SARS-CoV2 virus displayed visible infiltration of SARS-CoV2 cells in the bronchiolar epithelial boundaries, alveolar parenchyma, and renal tubular cells of kidney. Interestingly, thirty minutes of pretreatment with ACIS not only downregulated the expression of ACE-2, but also strongly mitigated the entry and cellular load of SARS-CoV2 in bronchiolar epithelial cells analyzed twenty-four hours post inoculation of virus. In fact, twenty-four hours treatment of SARS-CoV2 caused the thickening alveolar parenchyma with increased accumulation of mononuclear cells primarily inflammatory monocytes and macrophages. These are the indices of the onset of lung inflammation. Interestingly, KEPTIDE treatment significantly prevented the swelling and infiltration of inflammatory cells in small airway epithelium. Partial to significant degeneration of bronchiolar epithelial cells was also observed in virus-treated group. Surprisingly, KEPTIDE-treatment prevented the degeneration of these epithelial cells in virus-infected K18-hACE2 mice. We also observed increased vacuolization in kidney cortex with glomerular swelling after twenty-four hours of virus treatment, which is frequently observed in degenerative kidney [19, 20]. ACIS KEPTIDE significantly protected the kidney parenchyma with decreased vacuolization and also protected glomerular swelling.

### The infective property of SARS-CoV2 is not required for the death of COVID-19-infected animals

One most important highlight of this paper is to identify a new mechanism of SARS-CoV2-mediated death, which is entirely independent of its infective property. In our current project, we adopted βPL-mediated attenuation of infectivity in virus. This process strongly minimizes the risk of infection in users without compromising the purpose of our experiment, which is to measure the interaction and entry of the virus through ACE-2 receptor. These processes require the preservation of the 3D-structure of surface S-glycoprotein, that was not altered by βPL-treatment. SARS-CoV2 virions were thoroughly inactivated with 0.05% βPL-treatment. for 18 hrs in 4°C. After 18 hrs, the residual activity of βPL was neutralized with additional incubation at 37°C for 2 hrs. That process completely attenuated the infective property of virus. In accordance with the previous finding [13], our CPE experiment demonstrated that after βPL-treatment, virus completely lost its infective property. Interestingly, intranasal inoculation of that inactivated virus (1 *10^6^) significantly caused mortality in 28% of treated animals (2 out of 7) just after 24 hrs. The death was observed only in female mice. In addition to that, we observed reduced body weight, lowered body temperature, dropped oxygen saturation, and decreased heart rate in surviving animals. This dramatic and unexpected toxic effect after 24 hrs of inoculation with inactivated virus offered a paradigm shift in the molecular action of virus. Bpl-treated Inactivated virus does not have abilities to infect, replicate and spread, even though our results have demonstrated that it can cause substantial toxicity. Hence, our study identifies that the binding and modulation of ACE-2 receptor with spike protein seem to be the most critical step of SARS-CoV2-mediated death and toxicity. Upon binding, SARS-CoV2 might alter the physiological activity of ACE-2 receptor that potentially cause cytotoxicity and death. Our current report suggests that SARS-CoV2 does not require its traditional infective property to cause death.

### The peptide backbone of ACIS KEPTIDE cannot display similar protection

We compared the effects of KEPTIDE and its peptide backbone in improving lung and kidney pathologies in virus-infected K18-hACE2 mice. Several findings in this paper demonstrated that ACIS KEPTIDE™ displayed much stronger effect than its peptide backbone in terms of amelioration of lung and kidney pathologies, improving vital parameters of health, and preventing acute death response. In our previous report [8], we justified the significance of biochemical modification in the core peptide backbone of ACIS. Our cell-free assays demonstrated that these modifications are critical in order to enhance its binding affinity to ACE-2, physiological longevity, and robust absorption through intranasal path. Combining our previous findings with current results, we conclude that biochemical modifications of ACIS peptide backbone has significantly enhanced its protective effect in preventing SARS-CoV2-mediated toxicities.

### ACIS can be a vaccine alternative that exhibits prophylactic action against SARS-CoV2 infection

Several lines of our paper demonstrated that ACIS KEPTIDE could be a vaccine alternative. *First*, the pre-treatment with 50 μg/Kg Bwt of ACIS KEPTIDE for thirty minutes significantly protected the acute toxicity by SARS-CoV2. Surprisingly, KEPTIDE treatment not only protected the loss of body weight, temperature, heart rat, and oxygen saturation; but also prevented mortality. *Second*, the pre-treatment with the peptide backbone of ACIS KEPTIDE was found to worsen the vital signs of health in SARS-CoV2 infected animals. Although, no mortality was observed in peptide-treated group, 50% mice were moribund with severe motor impairment. *Third*, our immunohistochemical analyses of lung tissue revealed that the pre-treatment of ACIS KEPTIDE significantly prevented virus-driven histopathological alterations such as swelling of alveolar walls, integrity of bronchiolar epithelial membrane and hemorrhagic response in pulmonary vessels. Fourth, pre-treatment of ACIS KEPTIDE was also observed to protect glomerular integrity with decreased vacuolization in kidney cortices in SARS-CoV2-treated K18-hACE2 mice.

In summary, our current manuscript highlights the preventive role of ACIS KEPTIDE in the protection of acute toxicity and death caused by SARS-CoV2 virus and also demonstrates a novel mode of toxicity of SARS-CoV2, which does not require its genetic mechanism to be functional.

## Materials and Methods

### Acquisition and Handling of Wuhan standard SARS-CoV2

Wuhan standard SARS-CoV-2 (2019-nCoV/USA_WA1/2020) was kindly provided to KK by World Reference Center for Emerging Viruses and Arboviruses (TX, USA). Coppe Healthcare solutions has retained the approval, license and CDC-certification to handle SARS-CoV2 virus. Handling of viruses was performed in accordance with CDC-approved guideline. To generate virus stocks, Vero E6 cells were cultured in complete DMEM (+ 2% FCS), inoculated with virus at a MOI of 0.05. After 72 hrs, the virus-containing media was harvested, aliquoted, and kept at -80°C. Virus stock was titrated with a routine plaque assay on Vero E6 cells as described previously [8, 21].

### Virus inactivation and access to BSL-3 lab

SARS-CoV2 virus is infectious virus that can cause severe respiratory stress and pneumonia if not inactivated. Our aim is to explore if KEPTIDE blocks the entry of virus in lung cells. To perform this entry assay, the virus need not be in its infective state. Additionally, the inactivation also minimizes the health risk of users. Therefore, before virus administration, we performed the inactivation assay. Briefly, viral cells will be treated with 0.05% βPL for 18 hrs in 4°C. After 18 hrs, the cells will be kept at 37°C for 2 hrs for complete hydrolysis of βPL. That process will completely attenuate the infective property of virus and also completely nullifies the risk of handling SARS-CoV2 virus. This protocol has been optimized and reproduced by different scientific groups including KK in her CDC-approved CLEA-certified laboratory. The entire procedure of virus inactivation was performed in BSL-3 laboratory at Coppe laboratories. Coppe Lab has a dedicated Biosafety level 3 laboratory suite with one dedicated hoods, centrifuge, cell culture microscope, hot-water bath, two incubators, and a sink built in a secured containment room. Passage to that lab has entry through two sets of doors from access corridors. A clothing change room (i.e. anteroom) was located in the passageway between the two self-closing doors.

### Acquisition, justification, treatment, and disposal of K18-hACE2 mice

Six weeks-old K18-hACE2 mice (B6. Cg-Tg(K18-ACE2)2Prlmn/J) were purchased from Jackson laboratories (Stock #034860). Mice were housed in ventilated micro-isolator cages in an environmentally controlled vivarium (7:00 A.M./7:00P.M. light cycle; temperature maintained at 21-23°C; humidity 35-55%). Animals are provided standard mouse chow and water *ad libitum* and closely monitored for health and overall well-being daily by the investigator and supervised by a certified veterinarian in accordance with standard animal care guidelines. Before treatment, animals were transported to Coppe Laboratories in sealed cages covered with drapes as explained in our approved animal handling protocol. The entire virus inoculation work, perfusion and tissue collection were performed in a negative-pressure room inside BSL-3 hood. The justification of including eight animals per groups was derived with the following equation

***N* = [*z*^2^ × *p*(1 − *p*]]÷ *ε*^2^** = [1.28^2^ × 0.99 (1-0.99)]]÷ 0.05^2^ = **7. N** = sample size; **Z** = the z score, which is 1.28 for power 0.8; **p** = population proportion. For 98% confidence interval, p will be 0.98 & **(1-p)** = 0.02; and ***ε*** is the margin of error =0.05. For unexpected outcome of the experiment, additional 1 animal was ^included^ per group.

Once inactivated, these virus particles (1*10^6^; stock conc = 2.5*10^7^particles/ mL) were stored at 4°C overnight. On the day of experiment, mice were anesthetized with ketamine (50 mg/kg) and xylazine (5-7 mg/kg) mixture followed by the treatment with 50 μg/Kg Bwt KEPTIDE or peptide through intranasal route. After 30 minutes, we performed inoculation of 1*10^6^ virus in the nasal cavity. The virus stock was loaded in a micropipette and then gently disposed in both the nostrils by micropipette. After 24 hrs, animals were recorded for their vital health signs and then perfused with PBS followed by 4% PFA.

The removal of biohazard waste product was performed using proper institutional safety procedures. The perfusion waste was collected in a container that was thoroughly soaked with bleach and half-filled with concentrated bleach solution. For disposal of animal carcass, BSL-3 containment is strictly adhered to with proper bleach sanitization, locked in bleach-soaked biohazard bag, sealed with sealing threads, followed by placing into a potentially infectious material red waste container in the dedicated morgue. Final removal was performed by SteriCycle bio-waste disposal services.

### Anatomy and Histopathological evaluations

The formalin-fixed lung and kidney tissues was embedded in paraffin, cut into 5 μm thick sections, stained with hematoxylin and eosin (H&E) method, and evaluated for the histopathological changes by certified pathologist in MacNeal Hospital of Loyola Medicine. Histopathological changes in lung tissue such as infiltration of monocytes in alveolar parenchyma, thickening of alveolar walls, hemorrhagic response in and around pulmonary vessels, changes in bronchiolar epithelia were carefully scrutinized. Changes in glomerular structure, vacuolization in kidney cortices were monitored while analyzing pathological changes in Kidney tissue. Results were confirmed after evaluating n = 5-8 animals per group.

### Dual Immunohistochemical analyses

Five micron tissue sections were cut from paraffin-embedded lung and kidney tissue, installed in charged slides, deparaffinized in xylene, and then rehydrated through an ethanol gradient (100%, 95%, 80%, and 70%). The rehydrated slides were blocked with 2% horse-serum followed by incubation with primary antibodies (Mouse anti-S-glycoprotein antibody, 1:200 dilution, Millipore, Cat # MAB5676; Rabbit anti-ACE2 antibody, 1:200 dilution, Abcam, Cat # ab272690) for 2 hrs. After that, slides were washed with TBS + Tween (0.05%) solution for three times, incubated with mouse DAB- and rabbit AP-polymers for 1 hr, washed with TBS-T (X 3), reacted with DAB and AP substrates for color development, and finally stained with hematoxylin for nuclei. Slides were mounted with aqueous mounting media to preserve red color developed after AP reaction. Slides were thoroughly dried under air-blower at 70-80°C and imaged in a LOMO phase-contrast light microscope with ToupView 3.7 software. The dual immunohistochemical (IHC) analyses was performed as described in manufacturer’s protocol (Double Stain IHC Kit: M&R on human tissue; DAB & AP/Red; Cat # ab210059; Abcam).

### Immunofluorescence analysis in Calu-3 Cells

Calu-3, human lung epithelial cells, were cultured in complete DMEM for 72 hrs to achieve 70-80% confluency. After that, cells were starved with serum-free media for 1 hr followed by the treatment with 25 μM KEPTIDE™ for different time points such as 30 mins, 60 mins, 2 hrs and 6 hrs. After each time, cells were fixed with 4% PFA overnight. Next day, cells were thoroughly washed with PBST, incubated with Rabbit-anti ACE2 antibody (1:200 dilution) for 2 hrs, washed with PBST (X 3), incubated with 2° antibody [Alexa Fluor® 488 AffiniPure Goat Anti-Rabbit IgG (H+L); 1:200 dilution] for 1 hr, washed, and then mounted with fluoromount media. Our KEPTIDE tag shows autofluorescence at UV wavelength of 380-400 nm that enabled us to image slide in a dual channel under Olympus BX51 fluorescence microscope.

### Data collection, audiovisual recordings, and statistical analysis

Six to eight weeks old K18-hACE2 mice (*n*=8; 4 males and 4 females) were included to explore the toxic effect of KEPTIDE™ treatment. Mice were administered intranasally with 50 μg/Kg bwt KEPTIDE every day for 10 days and health vitals were measured as described previously [8]. Animals were tail-marked with marker pen before the treatment. Data of body weight, body temperature, heart rate, and O2 saturation were recorded in the notebook with date. Data were plotted in GraphPad Prism 8 software as XY scatter plots with X axis of “days” and Y axis of health variables.

Similarly, health parameters were recorded in notebook with dates after twenty-four hours of SARS-CoV2 treatment and also in KEPTIDE- and peptide-treated SARS-CoV2-inoculated in animals. Total 32 animals were shared in four different groups (vehicle only, virus only, virus + KEPTIDE, and virus + Peptide) with 8 animals per group.

For video recording, iPhone 7 camera was used. The cage was kept inside BSL-3 hood all the time with constant suction of air. BSL-3 room itself was located in a negative-pressure room. Users had performed the recording after wearing the full PPE including, gown, masks, two-layers of gloves holding cages far from the face. After video recording the phone was cleaned with bleach-soaked wipes.

Quantitative estimation of ACE-2-ir cells in bronchiolar epithelia and Mean color intensity of CE-2 expression in kidney tubular cells were performed in ImageJ software. Data were analyzed with a one-way ANOVA in a GraphPad Prism 8 software considering the treatment as a single factor. P-values were indicated with each figure to display the significance of mean between groups.

## Acknowledgements

The work is funded by the seed grant managed by JK and KKE. KK, GG and AR performed experiments; GG and AR analyzed data; AR designed research and wrote the manuscript. For business-related correspondence contact JK.

## Notes

### Competing Interest Statement

The authors have declared no competing interest.

