## Supplementary Fig for "Intranasal Administration of ACIS KEPTIDE™ Prevents SARS-CoV2-Induced Acute Toxicity in K18-hACE2 Humanized Mouse Model of COVID-19: A Mechanistic Insight for the Prophylactic Role of KEPTIDE™ in COVID-19"

Running title: Prophylactic Action of ACIS KEPTIDE™

To whom correspondence should be addressed:
Avik Roy, Ph.D.
SOTIRA LLC
Technology Innovation Center
10437 W Innovation Drive
Suite # 325
Wauwatosa, WI 53226

Ph: (773) 263 8182

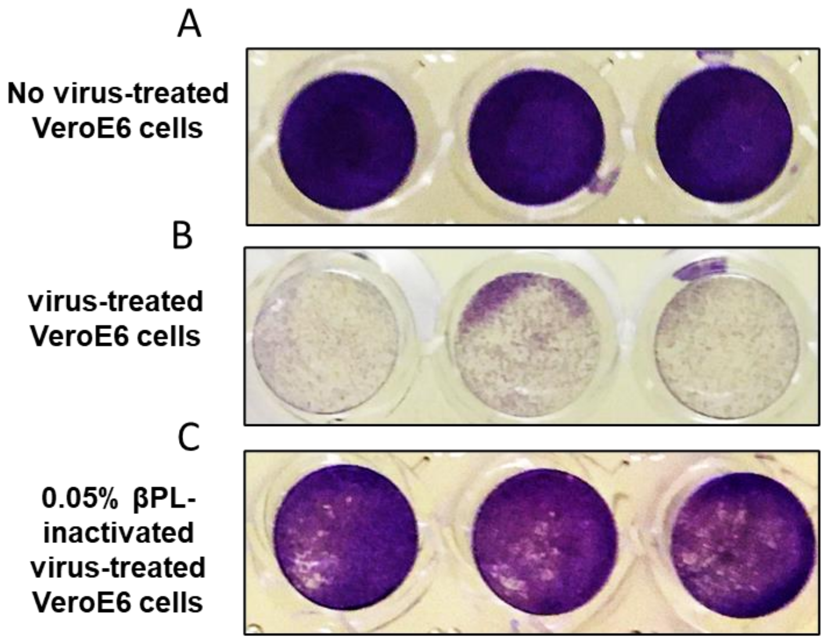


Supplementary Fig. 1

**Supplementary Fig.1. Inactivation of SARS-CoV2 virus with 0.05% Beta-propiolactone (βPL).** Viral stocks were treated with 0.05% βPL for 24 hrs at 4°C followed by neutralization of the residual βPL at 37°C. The viral stock was serially diluted and then plated on VEROE6 cells for 72 hrs followed by plaque assay to evaluate the inactivation. (**A**) No virus-treated VeroE6 cells with intact CV-stained agar monolayer. (**B**) Activated virus-treated VEROE6 cells with completely degenerated agar layer and (**C**) 0.05% βPL-inactivated virus with mostly intact agar layer.

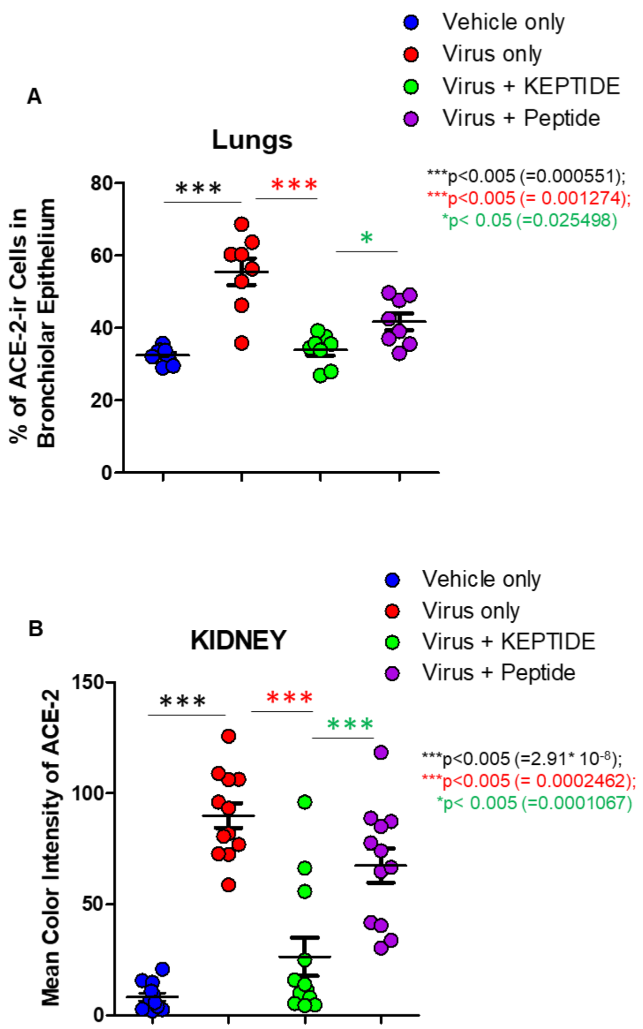


<u>Supplementary Fig. 2</u>

**Supplementary Fig.2. Analyses of ACE-2 expression in Lungs and Kidneys of Virus, Virus +KEPTIDE, and Virus + Peptide-treated K18-hACE2 animals.** (A) Percent of ACE-2 ir cells were counted as a function of eosin-stained cells in different groups. Total 9 images from three different animals were included in the counting (3 images from each animal). One way-ANOVA considering treatment as a single factor was performed to measure the significance of mean between groups, which results $F_{3,31}$ =20.33. (B) Men color intensity of ACE-2 ir cells were measured in renal tubular epithelium of different groups. Briefly, color channels were splitted in ImageJ, red channels were extracted, each epithelium was enclosed with a polygon tool, mean intensity was measured, and then subtracted with the blank. Total 12 images in three animals ( 4 images per animal) were included in the counting. One way-ANOVA considering treatment as a single factor was performed to measure the significance of mean between groups, which results $F_{3,47}$ =33.27. Results are mean ± SEM of three independent experiments.

**Supplementary video 1**

Inactivated SARS-CoV2-treated animals. After 24 hrs, significant mortality was observed with severe respiratory and motor impairment.

**Supplementary video 2**

Virus+KEPTIDE-treated group displayed significant protection in health and mortality.

**Supplementary video 3**

Virus+peptide-treated group displayed no mortality, but severe impairment in vital signs of health.
